# High sensitivity proteome-scale search for crosslinked peptides using CRIMP 2.0

**DOI:** 10.1101/2023.01.20.524983

**Authors:** D. Alex Crowder, Vladimir Sarpe, Bruno C. Amaral, Nicholas I. Brodie, Andrew R. M. Michael, David Schriemer

**Author notes:** Corresponding Author David C. Schriemer, Professor, Department of Biochemistry & Molecular Biology, Department of Chemistry, University of Calgary, 3330 Hospital Dr. NW, Calgary, Alberta, Canada. Authors contributed equally to the work (D. Alex Crowder, Vladimir Sarpe).

## Abstract

Crosslinking mass spectrometry (XL-MS) is a valuable technique for the generation of point-to-point distance measurements in protein space. Applications involving *in situ* chemical crosslinking have created the possibility of mapping whole protein interactomes with high spatial resolution. However, an XL-MS experiment carried out directly on cells requires highly efficient software that can detect crosslinked peptides with sensitivity and controlled error rates. Many algorithmic approaches invoke a filtering strategy designed to reduce the size of the database prior to mounting a search for crosslinks, but concern has been expressed over the possibility of reduced sensitivity with such strategies. Here we present a full upgrade to CRIMP, the crosslinking app in the Mass Spec Studio, which implements a new strategy for the detection of both component peptides in the MS^2^ spectrum. Using several published datasets, we demonstrate that this pre-searching method is sensitive and fast, permitting whole proteome searches on a conventional desktop computer for both cleavable and noncleavable crosslinkers. We introduce a new strategy for scoring crosslinks, adapted from computer vision algorithms, that properly resolves conflicting XL hits from other crosslinking reaction products, and we present a method for enhancing the detection of protein-protein interactions that relies upon compositional data.

## Introduction

Discovering molecular interactions in cells is an essential activity in proteomics and it has been the domain of affinity-based techniques for many years^1,2^. However, data returned from such techniques only indirectly identify binding partners. Direct interactions are in fact very difficult to detect, particularly in large-scale applications. For example, popular reporter-based optical methods like BRET (bioluminescence resonance energy transfer) can confirm pairwise interactions in live cells but require genetically tagging two proteins with sensors^3^. Useful real-time data can be returned on interaction behavior, but such pairwise methods are cumbersome tools for discovery.

Crosslinking mass spectrometry (XL-MS) is one technique that can provide a direct distance measurement between two points in proteome space. The basic approach involves chemically installing a bifunctional reagent between two reactive amino acid residues and detecting the linkage points using methods derived from bottom-up proteomics^4,5^. XL-MS can be used to model protein structure^6–8^, validate local interactome organization when paired with affinity isolation^9–11^, and even the map entire cellular networks^12–14^. Although the major elements of the technique have a long history, its application to whole cells for structural analysis is relatively recent, enabled primarily by higher sensitivity mass spectrometers and computational methods that can locate these direct linkage sites in highly complex digests.

The various applications of XL-MS present distinct methodological and computational challenges. For example, structure determination requires extensive crosslinking to produce enough constraints for accurate model building. Such an exercise has a moderate tolerance for false positives, provided that the crosslinks sample the structure in an unbiased manner^15,16^. On the other hand, defining an *in situ* protein network could be achieved with as little as one crosslink per interaction. In this case there is a very low tolerance for false positives^17^. A growing number of computational tools have been developed to meet the challenges of crosslink identification for these applications^18–20^, but we need new approaches as tool comparisons reveal performance gaps^18,19,21,22^.

Specifically, a major unmet need involves the efficient detection of crosslinks in LC-MS/MS data obtained from ultra-complex mixtures, particularly whole proteomes^23,24^. There are solutions, but they remain difficult to scale and are focused on a narrow selection of crosslinking reagent types. Some of the earliest computational methods were developed to support searches of limited complexity, rarely involving more than a few 10’ s of proteins^25,26^. The fundamental challenge to scaling these early approaches involved the combinatorial nature of the search space. Methods that evaluate all possible peptide couplings, restricted only by the precursor mass of the crosslinked peptides, struggle with whole proteome searching. Even very strict peak matching criteria and efficient data structures^24^ remain inefficient because the search scales as n^2^ (where n is the number of peptides in a search library). One elegant solution to this problem is an experimental one. Gas-phase cleavable crosslinkers such as PIR, DSSO and DSBU generate linear peptides in MS^2^. These can be selected for peptide fragmentation in MS^3^ to facilitate a more straightforward proteomics search^27,28^. Unfortunately, MS^3^-based detection methods present sensitivity and duty cycle issues^29^. An MS^2^-based alternative using stepped HCD fragmentation is an effective work-around^29,30^ but much computational efficiency is lost in defaulting to MS2 analyses. The challenge of cleavable reagents appears to dominate software development for whole proteome searching. However, there are many noncleavable reagents that can tune crosslink yield and distance measurements, plus they can provide cross-peptide fragment ions that are very useful in validating IDs. Thus, while we require more tools for XL-MS data analysis in general, the need for noncleavable reagents is particularly great.

Any computational solution that addresses the *O(n*^*2*^) time complexity problem should not trade sensitivity and accuracy for computational efficiency. There is little to be gained from developing a fast solution that either misses large numbers of crosslinked peptides or has an unacceptably high false positive rate. Several approaches to the problem have been considered. Strategies include accelerated implementations of the brute-force method described above, but more commonly, software packages use a two-pass search in which a method is first applied to restrict the search space, followed by a more refined scoring strategy that acts on the reduced database^21,31,32^. There are many ways to implement a two-pass method. For gas-phase cleavable crosslinkers, the MS^2^-liberated peptides provide peptide masses that can help narrow the search space^33^. More typically for all crosslink types, database reduction uses an open-modification search that was inspired by routines for detecting unknown post-translational modifications^34^.

The exercise is defined as a search for a peptide with a modification of unknown mass, that is, the linked second peptide. Each peptide receives a preliminary score. Candidate peptides with scores above some threshold are selected and ranked, then used to restrict the search for the second peptide^21,31^. The resulting combinations are rescored in a full search against the MS^2^ spectrum. Variations of the two-pass approach exist, usually differing in how the precursor mass is used to filter candidates. However it has been suggested that a presearch strategy may not advance sufficiently high numbers of peptides to be sensitive enough in proteome-scale applications^24^. This would be unfortunate as brute-force methods may never be practical for proteome-wide searches with modest computational effort, especially if post-translational modifications need to be considered.

We updated our CRIMP crosslink detection software^35^ with a new library reduction strategy to determine if whole proteome searches can be sensitive, fast and independent of the style of crosslinker used. The first version of CRIMP used a prescoring approach that required sparse evidence for the existence of *both* linear peptides in the MS^2^ spectrum, in the form of fragment fingerprints. The approach was surprisingly sensitive, given that only the free forms of the peptides were searched in the first pass, but design limitations prevented extending the concept to whole proteomes. CRIMP 2.0 incorporates an improved library reduction engine and a new scoring algorithm that resolves spectral conflicts across all categories of hits (*e*.*g*., free peptides, monolinks and crosslinks). Our revised approach to error estimation accounts for cross-category spectral conflicts, mostly ignored in other search tools, and supports a new method of detecting of protein-protein interactions. We demonstrate that a library reduction strategy can indeed deliver high sensitivity and support whole proteome analysis of noncleavable and cleavable experiment types, with modest computational resources.

## Experimental Section

### Synthetic peptide benchmark dataset 1, with variant

Data from a synthetic crosslinked peptide library was accessed from PRIDE (PXD014337), which contains a set of crosslinked and monolinked peptides engineered from amino acid sequences of 95 tryptic peptides derived from *S. pyogenes* Cas9^36^. These peptides range in length from 5-20 residues and contain only one linkable lysine. They were prepared in distinct groupings to aid in false positive identifications and contained a maximum of 426 crosslinked peptides. Data from the DSS crosslink library were used and searched in several ways. First, the data were searched against a database consisting of Cas9 and various other databases for entrapment, including 116 proteins from the cRAPome^37^, the *E*.*coli* proteome (UP000000625, 4448 entries), the *S. cerevisiae* proteome (UP000002311, 6060 entries), the *D. melanogaster* proteome (UP000000803, 22072 entries), and the *H. sapiens* proteome (UP000005640, 79759 entries). Second, to simulate interprotein crosslinks the same databases were searched, but Cas9 was first segmented into four distinct subunits in such a manner that subunits A, B, C, and D could form both intra and inter-protein crosslinks. Here, inter-protein crosslinks are the most abundant: 248 potential inter-links and 178 potential intra-links (which here includes homotypic crosslinks). This segmentation, and the parameters in the CRIMP search, can be found in Supporting Information. In the first strategy, crosslinks were considered correct if crosslinked peptides were assigned to the right peptide group, simulating an intraprotein crosslink search. In the second strategy, crosslinks were considered correct if they were assigned to the right peptide group and the right inter-subunit interaction, simulating an inter-protein crosslink search.

### Synthetic peptide benchmark dataset 2

Data from a second synthetic crosslinked peptide library was accessed from PRIDE (PDX029252), which contains a set of crosslinked and monolinked peptides engineered from amino acid sequences of 141 peptides from 38 proteins of the *E*.*coli* ribosomal complex that more intentionally allows for testing of inter- and intra-FDR^30^. Data from the DSSO crosslinked libraries were used, with varying degrees of added peptide noise and database entrapment as per the published study. The sets contain a maximum of 1018 crosslinked peptides. The parameters in the CRIMP search can also be found in Supporting Information.

### Proteome-scale *E.coli* dataset

Data from a large crosslinking experiment was accessed from the ProteomeXchange Consortium partner repository jPOSTrepo under the accession codes JPST000834 and PXD019120^38^. In this study, *E. coli* K12 strain was produced and lysed, followed by separation of the soluble high molecular weight proteome by size exclusion chromatography. Proteomics experiments on each SEC fraction identified the proteins that comprise potential interactions within a given fraction. Each fraction was then crosslinked two ways: with bis(sulfosuccinimidyl suberate) (BS3) and with disuccinimidyl sulfoxide (DSSO). For each crosslinker, fractions were then pooled and digested with LysC/trypsin followed by a two-dimensional separation of the resulting peptides. Briefly, each digested fraction was separated using a PolySulfoethyl A SCX column, with each fraction of this separation then separated by hSAX. Each crosslinking experiment generated 90 fractions and two replicates (nominally 180 LC-MS/MS runs). Database searches incorporated the approximately half of the *E. coli* proteome, as per Rappsilber *et al*.^38^ The parameters used in the CRIMP searches for each crosslinker can be found in Supporting Information. Crosslinks were considered correct if crosslinked peptides involved proteins detectable above a specified intensity threshold for a given SEC fraction^38^. A resulting master list of 544,274 plausible PPI’ s was mined for error estimation, along with 8,914,801 interactions that were deemed non-crosslinkable. FDR-based error estimates were adjusted for the size of the false and plausible search spaces.

### Software design and availability

CRIMP2.0 was built within the Mass Spec Studio design framework, a repository of data analysis routines available within the Studio for structural mass spectrometry applications such as HDX-MS, covalent labeling, and integrative structure modeling. Software was written in C# and it leverages an extensive repository of reusable content. All the processed results can be regenerated using the datasets listed above, using CRIMP2.0 from www.msstudio.ca (version 2.4.0.3544.

## Results and Discussion

### Strategies for library reduction

All current concepts for crosslink detection use one of two methods to reduce the search space (**Figure 1**). The brute-force enumeration of all peptide pairs constrains the search through the precursor mass, by determining the set of all possible α and β peptide masses that sum to the precursor mass, after accounting for the addition of the linker itself. All relevant combinations are then scored directly against the MS^2^ spectrum of the crosslink peptide in a 1-pass determination. Precursor mass is a relatively weak constraint, particularly when the mass tolerance is relaxed, and is the primary source of the search’ s high time burden. Large numbers of combinations result when scoring the whole proteome. In return for this weak constraint, the overall search has the potential to be very sensitive as in theory no combinations will be missed.

**Figure 1.**
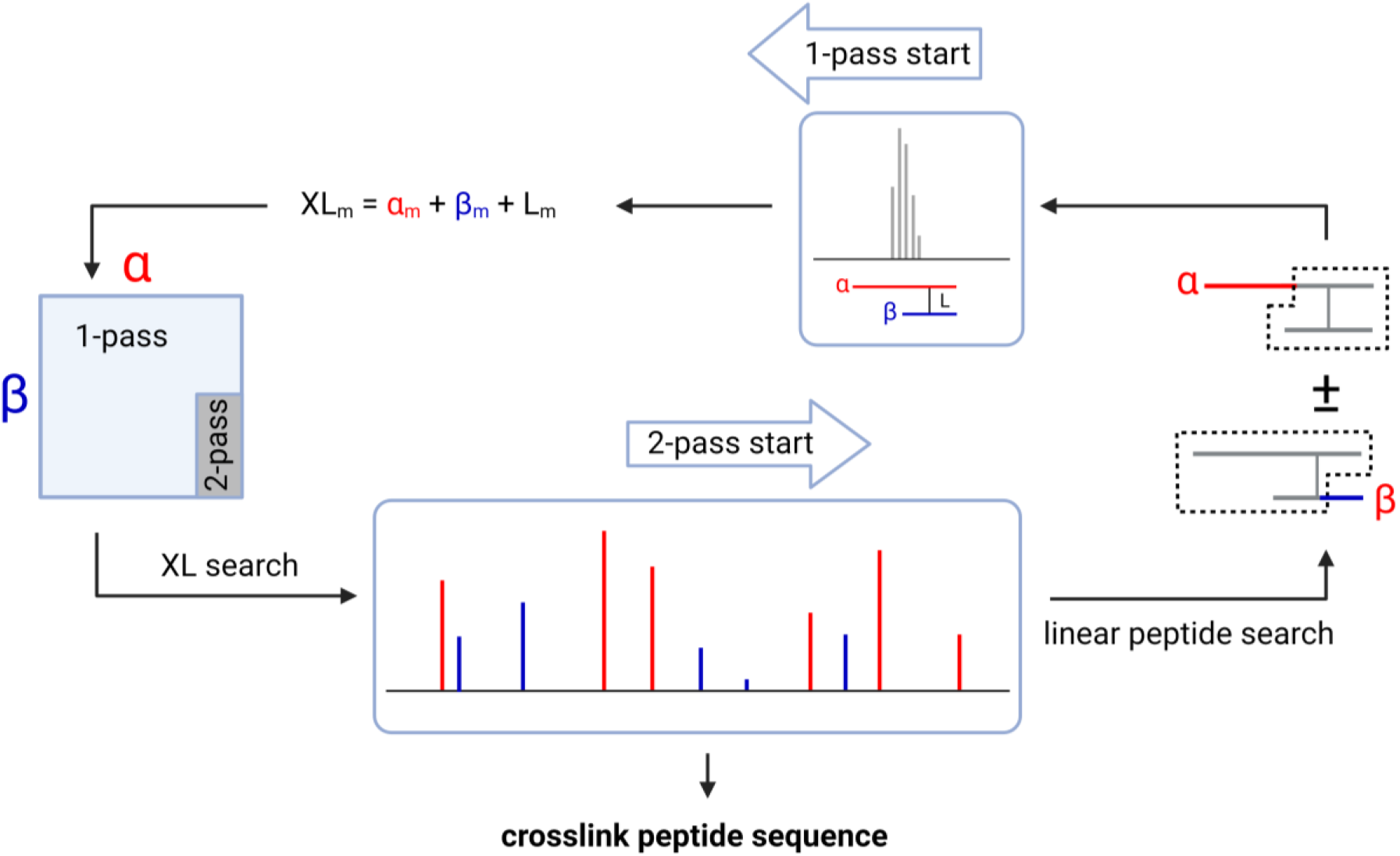
Schematic outlining the typical approaches to searching MS^2^ datasets for evidence of peptide crosslinking. The 1-pass approach begins with the precursor mass of the putative crosslinked peptide and constrains a database search through a simple three-term sum involving the α peptide mass, the β peptide mass and the linker mass. Combinations are then searched against the MS^2^ spectrum. The 2-pass method begins and ends with the MS^2^ spectrum. First, candidate α and β peptides are found in the MS^2^ data, and only then is precursor mass used to constrain combinations for a more exhaustive search of the MS^2^ data.

The other concept invokes a 2-pass approach, in that the MS^2^ spectrum of the crosslink peptide is mined twice. The first pass attempts to identify options for the dominant α peptide through a linearized fragment set, a subset of the total fragments in the spectrum. That is, a database search looks for fragments arising from the free peptide plus its mass-modified form. Detecting the mass-modified form requires an open modification algorithm that treats the second peptide (plus linker) as a modification whose mass can be calculated by subtracting the candidate α peptide mass from the precursor mass. In this fashion, the α peptide sequence can be read past the crosslink site. This open modification search is also dependent upon the precursor mass, but the selection of α peptide candidates is primarily driven by the quality of the MS^2^ hits. Most search tools restrict this presearch of the MS^2^ spectrum to the α peptide alone^21,31^, relying again on the precursor ion to define the candidate β peptides by subtraction of the α peptides. These tools avoid mounting a linearization of the β peptide, arguing that the MS^2^ spectral data for the β peptide is comparatively sparse because of preferential fragmentation^39^. Nevertheless, linearizing both peptides has been successfully demonstrated in the Goodlett lab^40^. As well, the original Kojak search tool looks for both and then constructs the peptide combinations for rescoring^41^, but it underperforms those tools that restrict reduction to the α peptide, which seemingly validates the concerns expressed over the value of the β peptide sequence information.

We wanted to explore the double peptide presearch concept further, as search performance can be affected by many elements of the workflow. This exploration was inspired by the demands of the *in situ* crosslinking application itself. In whole proteome searching, particularly where the goal is to identify protein-protein interactions from as little as one crosslink match, we argue that both peptides must generate fragment series of sufficient quality to identify two proteins or protein groups. A search tool that finds poor-quality β peptides is of little value, especially if it reduces computational efficiency. To minimize the computational limitations associated with precursor mass constraints, the original version of CRIMP explored a pure linear peptide search for both α and β peptides in a 2-pass method. This returned a sensitive search, somewhat surprisingly given the incompleteness of these fragment subsets^42^. Including peptide linearization for both α and β peptides further increased crosslink detection sensitivity in preliminary testing (not shown), suggesting to us that this library reduction method may be superior to a strictly α-based strategy for rapid whole proteome searching.

### Upgrading crosslink analysis in the Mass Spec Studio: CRIMP 2.0

We incorporated this reduction strategy into a revised search routine that was designed to support crosslink identification for all applications – from structure modeling to whole proteome searching (**Figure 2**). Improvements were made to crosslink rescoring and error estimation, supported by improvements in signal processing. Several computational enhancements were implemented to accelerate search speed and improve the navigation and export of search results. These are described briefly below.

**Figure 2.**
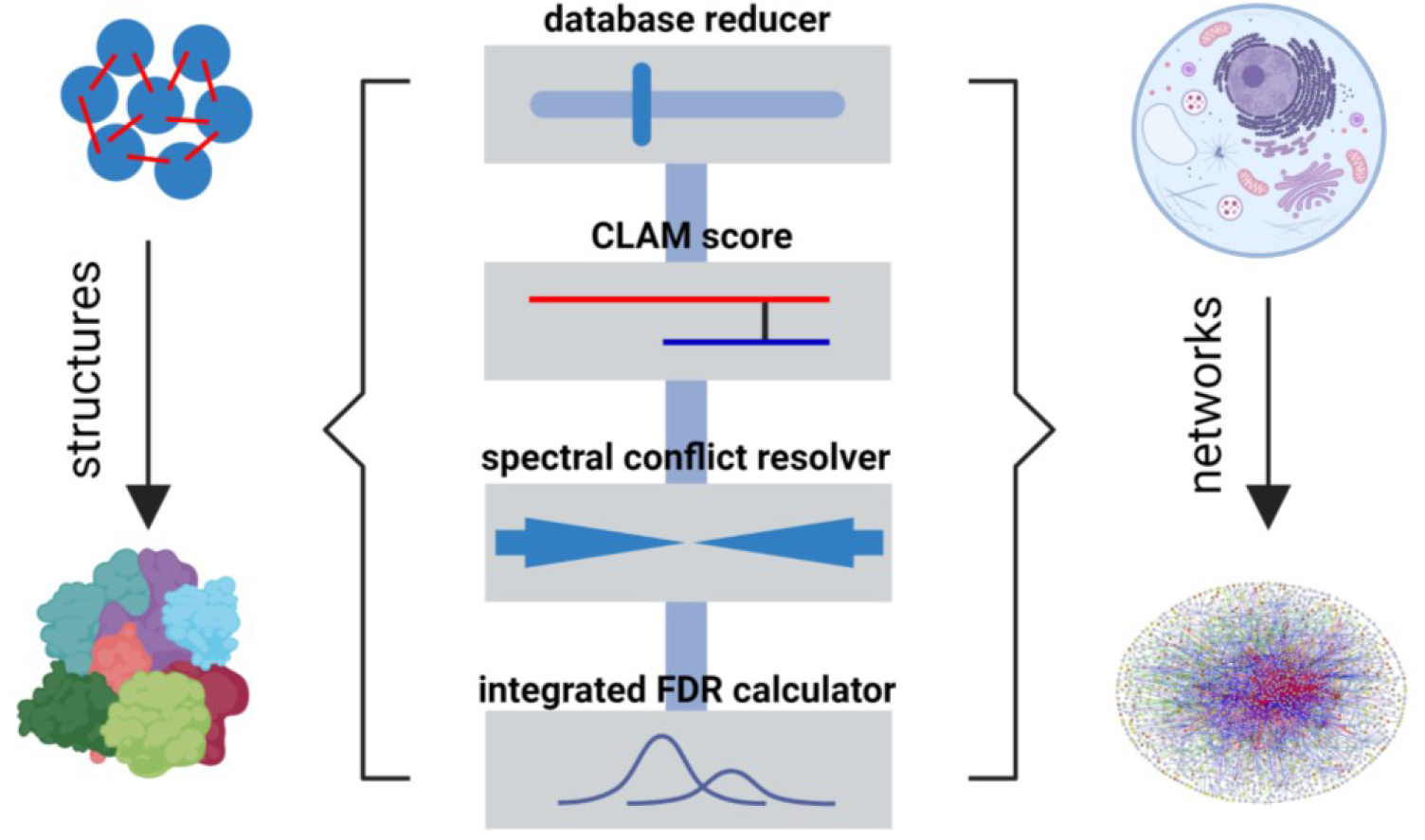
Key elements of the CRIMP 2.0 functionality. Database searches can be tailored to the XL-MS application with a minimum of parameterization. Search setup and results navigation are supported in a user-friendly package.

#### Spectral annotation and fragment assignment

Scoring is supported with several improvements to signal processing. These include an adaptable method of precursor ion detection that is sensitive to the S/N of the given feature. The goodness-of-fit of an isotopic distribution for a candidate peptide can be relaxed when the S/N is poorer, allowing the natural advantages of tandem mass spectrometry to override poor MS ion statistics. Improvements in the identification of monoisotopic peaks were adopted for the MS^2^ domain as well, to support both centroid and profile-mode datasets. A new approach to fragment ion assignment was also adopted. Fragment types are numerous and complex in crosslinked peptide spectra, and certain fragment types will draw from a much larger distribution of possible values than others. For example, the number of possible internal fragment ions for crosslinked peptides (*i*.*e*., generated by two cleavage events) is much larger than simple single cleavage events, but it is inappropriate to allow for equal weighting of these internal fragments during fragment assignment given that they are usually less abundant than other sequence ions. We developed and tuned a greedy assignment algorithm based on a hierarchy of fragment types, and the resulting identifications and weights are used in place of the number of detected peaks for crosslink scoring.

#### Scoring and conflict resolution

CRIMP uses a probabilistic scoring of α and β peptide candidates based on E values rather than a simple peak count to assemble a more informative ranked list of candidates. This is based on our OMSSA+ scoring method^35^, adapted for high-resolution and multiple charge state ions. The pre-search concept invokes a user-adjustable sensitivity setting for each peptide (the E-score). A scoring and ranking strategy was developed to ensure that high-probability candidate peptides are enriched within the top N groups of hits, where groups are defined as candidate peptides with similar scores. Top N is adjustable, but experimentation has shown that a value of 10 is sufficient in most every scenario, and the dependency on the scores for the α and β peptides is well-behaved (**Figure S1**). Peptides are then paired and exhaustively scored against the MS^2^ data.

Crosslinking reactions produce crosslinked peptides in relatively low abundance, and the reaction products contain comparatively high amounts of mono-linked, loop-linked and free peptides. Some of these can be present even after enrichment technologies are used. As a result, many annotated fragment peaks can be shared between these products, particularly when they share sequence. Resolving these conflicts is not always straightforward as these different products tend to generate unique scoring ranges and noise distributions. We developed a strategy that manages conflicts and normalizes scores across products to determine the best overall match to a spectrum. Briefly, a multi-term scoring vector inspired by computer vision algorithms was created that contains several OMSSA+ scores calculated from subsets of fragments identified in the MS^2^ spectrum. These subsets are selected based on their ability to discriminate between different digestion products. Scoring vectors are determined for all possible matches and a given identification is penalized using the shared fragment assignments from possible conflicts. This penalized scoring vector is transformed using q-values and then a final CLAM (Competitive Label Assignment Method) score is determined from the inner product of the transformed scoring vector (see Supporting Information).

#### False discovery rate (FDR) estimation and aggregation

We use FDR calculations to assess the total error in search results, involving decoy database searches to estimate the error. Database searches and FDR calculations are usually performed in a category-based manner and are not able to address the impact of cross-category misidentifications on FDR (where category refers to the various products of a crosslinking reaction). Additionally, there can be a sparse numbers of decoy hits for any given category, which makes FDR estimation less precise as a result. All categories should be integrated in a joint FDR calculation, but it requires a method for normalizing score distributions across categories. There is considerable variation in database size for the different categories and thus in the underlying search noise. For example, the search space for inter-protein crosslinks is considerably greater than the search space for intra-protein crosslinks, necessitating different decoy databases. Thus, to integrate all categories of data we standardize CLAM scores according to the specific error distributions of each, and then all categories of scores are combined. A scaling step transforms these scores to a uniform range of 1-100 to permit direct score comparisons between categories (*e*.*g*., mono-links and crosslinks). Only then is the final FDR estimate determined, and it is calculated as a geometric average of the global and the local FDR estimates as a concession to the validity of both approaches (see Supporting Information).

Search results are then aggregated within runs, and between runs in the case of replicates. The first step of the aggregation workflow is to reduce crosslink spectrum matches (CSMs) to unique peptides and peptide pairs, by collapsing redundancies arising from charge state, retention times and variable modifications. Next, higher level aggregations to unique residue pairs (URPs), proteins, and protein-protein interactions (PPIs) are made, with each level supported by a separate error calculation. However, PPIs are not sensitively determined with these crosslink-centric methods of detection and error estimation, likely because the “all by all” search space is an overestimation of the true PPI space and drives up the noise level of the search. We reasoned that PPIs should possess evidence for their component proteins in the form of free peptides, mono-links, loop-links and intra-protein links. To this end, we rescore PPIs before error estimation by calculating a transformed value that includes evidence of the presence of component proteins and the success of the labeling chemistry (see Supporting Information).

#### Computational enhancements

We also significantly improved the way searches are supported computationally, particularly in database reduction. A peptide fragment index approach is a useful way to streamline a search. It involves generating fragments for every peptide in a database, and then scoring the peptide against all MS^2^ spectra based on the fragments generated. The approach is used by MSFragger for whole proteome searching, and by pLink2 for α peptide searches^21,43^. We extended this fragment index concept to the whole protein level, so that every unique fragment from every protein is only scored once per spectrum during the library reduction step. The increase in computational efficiency is considerable, and the approach even limits database expansion when incorporating variable modifications and missed cleavages.

Many XL-MS applications require replicate runs, such as quantitative crosslinking or when rerunning samples to saturate identifications in data-dependent acquisition (DDA) experiments. To improve robustness in the generation of candidate α and β peptides, the reduced library is propagated between replicate searches in a manner that is reminiscent of “match between runs” in proteomics^44^. Here it means that we share the reduced library between replicates for any given precursor mass. Replicate DDA runs can generate some variation in crosslinker sampling, leading to unequal sets of candidate peptides during reduction. Propagation therefore combines the candidate lists of peptides across all replicates, followed by a precursor assignment pass using the complete set of peptides.

A set of changes was made to manage resources on more limited platforms like desktop and laptop computers. The user can adjust the degree of stringency on pre-searching α and β peptides (as well as top N), creating the opportunity to conduct a search that approximates a brute-force strategy for samples of limited complexity, or one that applies strong reduction for whole proteomes. This gives the user the potential to balance concerns over sensitivity with computational performance, however we will demonstrate below that the tradeoff is not a problematic one, and optimal searches converge on a very narrow set of parameters. Additional improvements include the aggregation of similar spectra prior to search and the use of vectorized computations wherever possible. Crosslinking datasets are also sliced into subsets of spectra and processed serially, to manage memory usage.

Finally, numerous improvements were introduced to support the full range of crosslinking experiments and it anticipates new crosslinker designs. The new version preserves the validation-based routines and spectral navigation tools of version 1, which are particularly useful in structure modeling applications of XL-MS (**Figure 3**). We include visualization and export utilities for all downstream applications. Cleavable crosslinkers are accommodated through an MS^2^-based workflow, as opposed to the less efficient MS^3^-based routine. Tool tips are included throughout, aided by an extensive glossary of terms.

**Figure 3.**
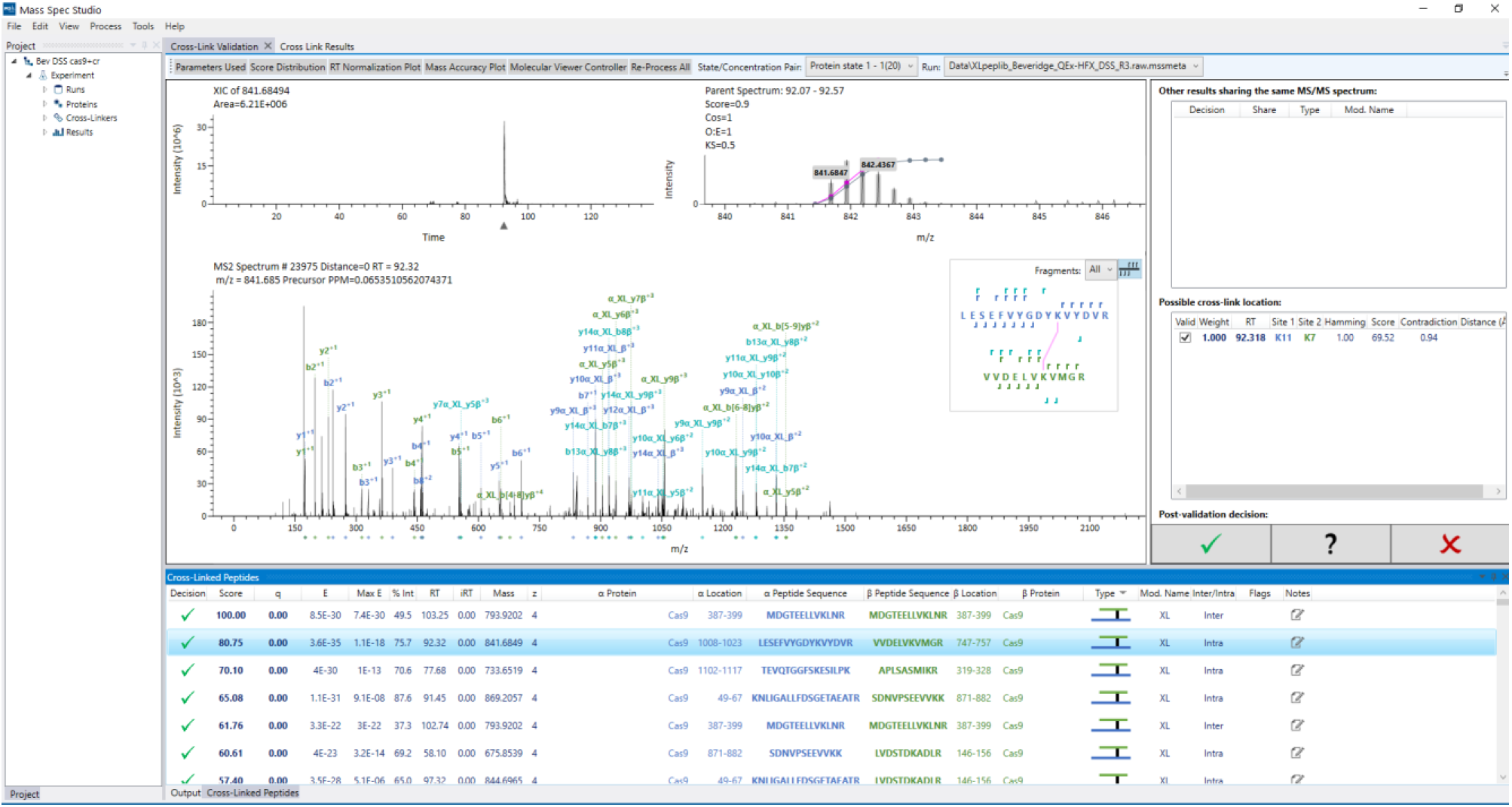
A screenshot of the post-search validation viewer in CRIMP 2.0, which provides rich functionality for navigating, validating, and exporting results.

### Evaluation of search sensitivity

We applied CRIMP 2.0 to database searches of increasing complexity and scale. First, we reanalyzed the triplicate datasets from the synthetically generated crosslinked peptides provided by Beveridge *et al*.^36^ For the DSS noncleavable linker set, we detected 226±7 unique crosslinks at a real FDR of 2.9% (for an expected FDR of 5%). The real FDR is calculated based on the known composition of the peptide set. Search sensitivity meets or exceeds several other tools that have operated on this dataset (**Figure S2**). We used an E_α_ peptide setting of 10% to reflect the higher quality of the dominant peptide and allowed the E_β_ setting to be 99%, strongly relaxing the requirement for a high-quality β peptide. These are the default values in CRIMP 2.0 for all searches.

To compare with a brute-force approach, CRIMP 2.0 identified 1341 CSMs with a real FDR of 2.0%, compared to 885 CSMs and a real FDR of 3.1% for OpenPepXL. In this example there is a slight positive effect of propagating the reduced libraries between runs, which disappears when the results are aggregated. The lower sensitivity of the brute force approach is likely related to the higher level of noise that accompanies a search of all possible pairwise combinations. We then searched the replicate noncleavable crosslinker data at two different FDR cutoffs (with database propagation) at various levels of database entrapment (**Figure 4A,B**). The sensitivity decreases, but approximately 35% of the detectable unique crosslinks are still found with almost 80,000 proteins as entrapment. To evaluate our approach for inter-protein detection, we altered the database from its original design by segmenting the Cas9 protein into four smaller “proteins” (see Experimental Section). This segmentation turns most intra-protein crosslinks into inter-protein crosslinks. The results show a diminishing sensitivity for inter-protein crosslinks at higher levels of entrapment, but the rate of reduction is only slightly greater than intra-protein crosslinks (**Figure 4C,D**). Error estimation begins to destabilize at higher entrapment, but this is due to the very low number of hits returned in the searches. These searches are computationally efficient even with significant levels of entrapment, highlighting the utility of a strong database reduction concept (**Figure S3**).

**Figure 4.**
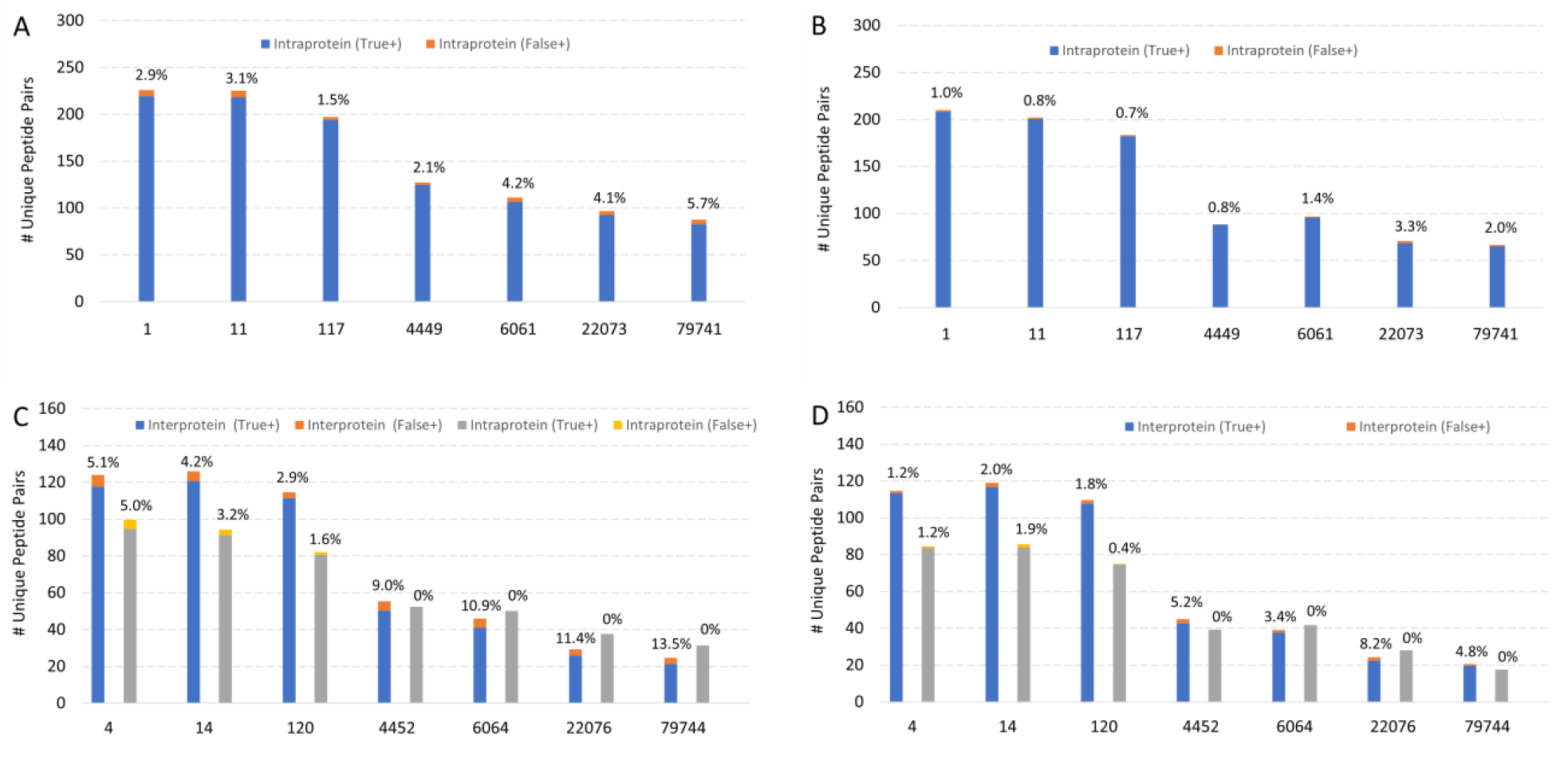
Crosslink sensitivity determination using the replicate dataset from Beveridge *et al*.^36^ for the DSS crosslinker. The analysis for the Cas9 database at (A) 5% and (B) 1% calculated FDR. The results for the segmented Cas9 database at (C) 5% and (D) 1% FDR, presented as both intraprotein and an interprotein search results. Effect of entrapment is shown using multiple added databases with the noted protein complexity.

We then expanded the tool comparison using a new set of replicate data published by Matzinger *et al*.,^30^ involving a significantly expanded set of synthetic crosslinked peptides. We specifically focused on cleavable crosslinkers and used our default search conditions. Database propagation was deactivated to promote a fair comparison with the original study. Again, CRIMP 2.0 outperformed all tools at both the expected and real FDRs (**Figure 5**), detecting 673 (66%) of all theoretically possible crosslinked peptides on average, and 760 (75%) when all replicates were aggregated after the search. Activating propagation had no effect on the results (not shown), likely due to the high abundance and quality of the crosslinked spectra. Propagation is anticipated to be impactful in lower abundance and/or under-sampled datasets. Repeating the searches at an expected FDR of 5% provides an indication of the value of propagating the reduced database, where 699 hits on average were identified without propagation and 730 with propagation. Under more typical search conditions where free peptides are present CRIMP 2.0 performed equally well, particularly at the high complexity searches with extensive entrapment (**Figure S4**). FDR estimation is reasonably accurate in these peptide-contaminated datasets (1-3.4%), even with extensive entrapment. Much larger datasets are likely required to generate a more precise estimate.

**Figure 5.**
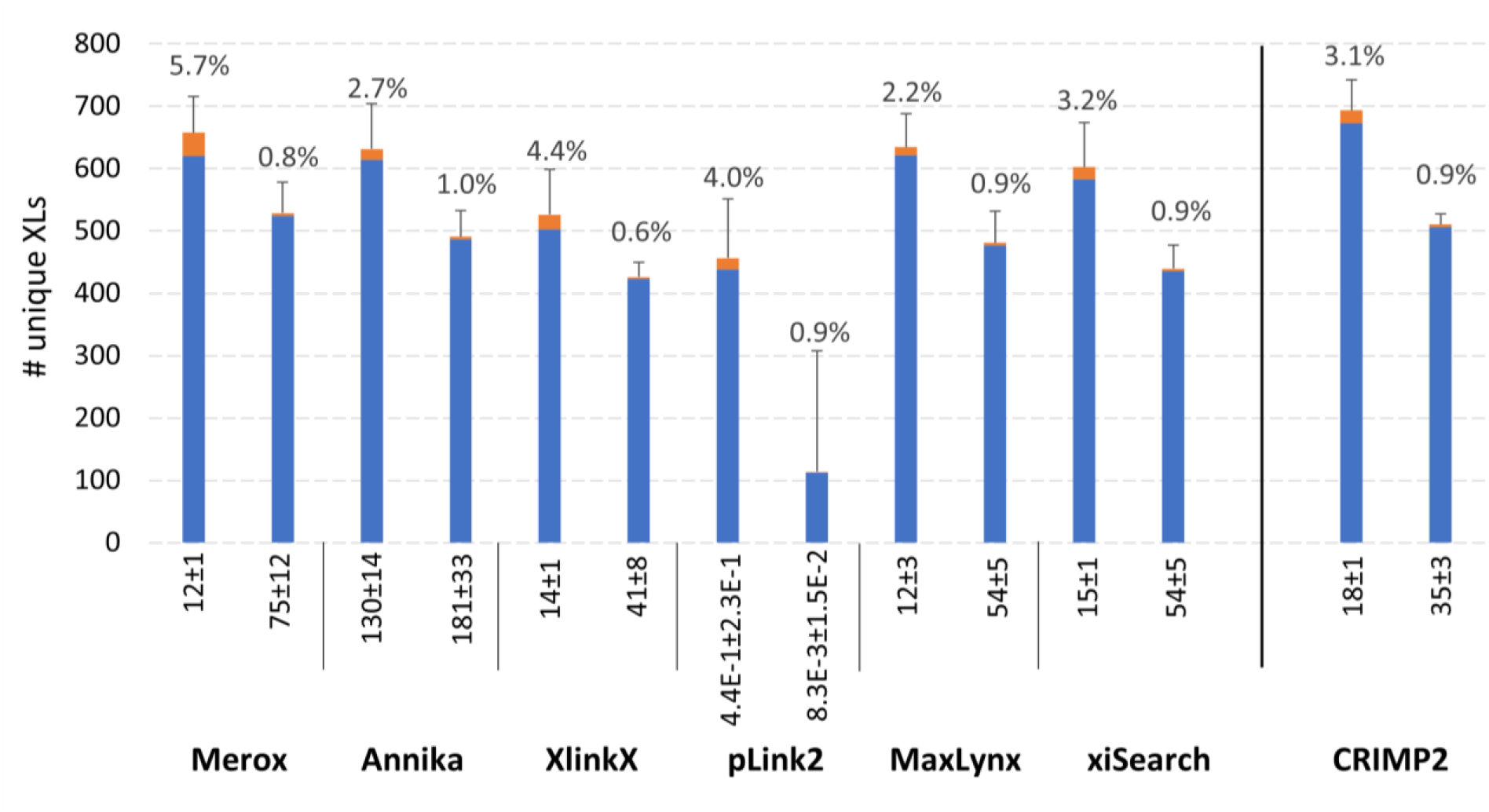
Average crosslinked peptide numbers using DSSO as the crosslinking reagent and a stepped HCD MS^2^ method for data acquisition. All results for the noted algorithms are derived from Matzinger *et al*.^30^ with the addition of CRIMP 2.0, at an expected FDR of 1% (left bar in pair), and corrected results using a post-score cutoff to reach an experimentally validated FDR of 1% (right bar in pair). True positives in blue, false positives in orange. Real FDR posted as callouts. Error bars indicate standard deviation of the total hits, n=3.

CRIMP 2.0 uses a new strategy for detecting PPIs. It has been the practice in the community to restrict a database search to only those proteins that are detected in the sample being crosslinked^38^. This detection is achieved with a separate proteomics analysis. The approach helps constrain the search space (and thus accelerate the search) and it also helps to limit the generation of false positive identifications. Other methods use some element of compositional analysis to restrict the peptide search space^45,46^. CRIMP’ s PPI scoring method incorporates all internal evidence for the existence of a given protein in the score but does not demand it (Supporting Information). That is, the detection of other crosslinks and all other possible reaction products for a given PPI are incorporated into the score. We and others have observed that *bona fide* PPIs often present multiple inter-peptide crosslinks, and thus should be granted a higher confidence identification, but single inter-protein crosslinks do occur and should not be penalized. Our scoring method is applied during the aggregation of peptide identifications and is designed to boost PPI scores but not override these high-quality individual inter-protein crosslinks. To demonstrate, we allocated all the possible crosslinked peptides from the synthetic peptide library of Matzinger *et al*.^30^ to their ribosomal proteins of origin. PPIs generated from searching the resulting library will not be physically meaningful, but it will still generate a searchable PPI space. This exercise generated 331 PPIs after removing redundancies, 300 of which are heterotypic and 31 that are homotypic. At 1% FDR, we detect 313 (95%) of the possible PPIs in a database search restricted to 171 proteins, which diminishes to 151 (46%) with heavy entrapment (20,334 human proteins) (**Figure 6**). These results indicate a robust scoring method. We see strong FDR control when searching crosslinks heavily “contaminated” linear peptide data, which shows that compositional data (in the form of free peptides at least) does not overwhelm the requirement for quality crosslink identification. Deactivating the compositional boost for this dataset has a negligible effect on the FDR and indicates the benefit to PPI sensitivity in adding compositional data (**Figure S5**).

**Figure 6.**
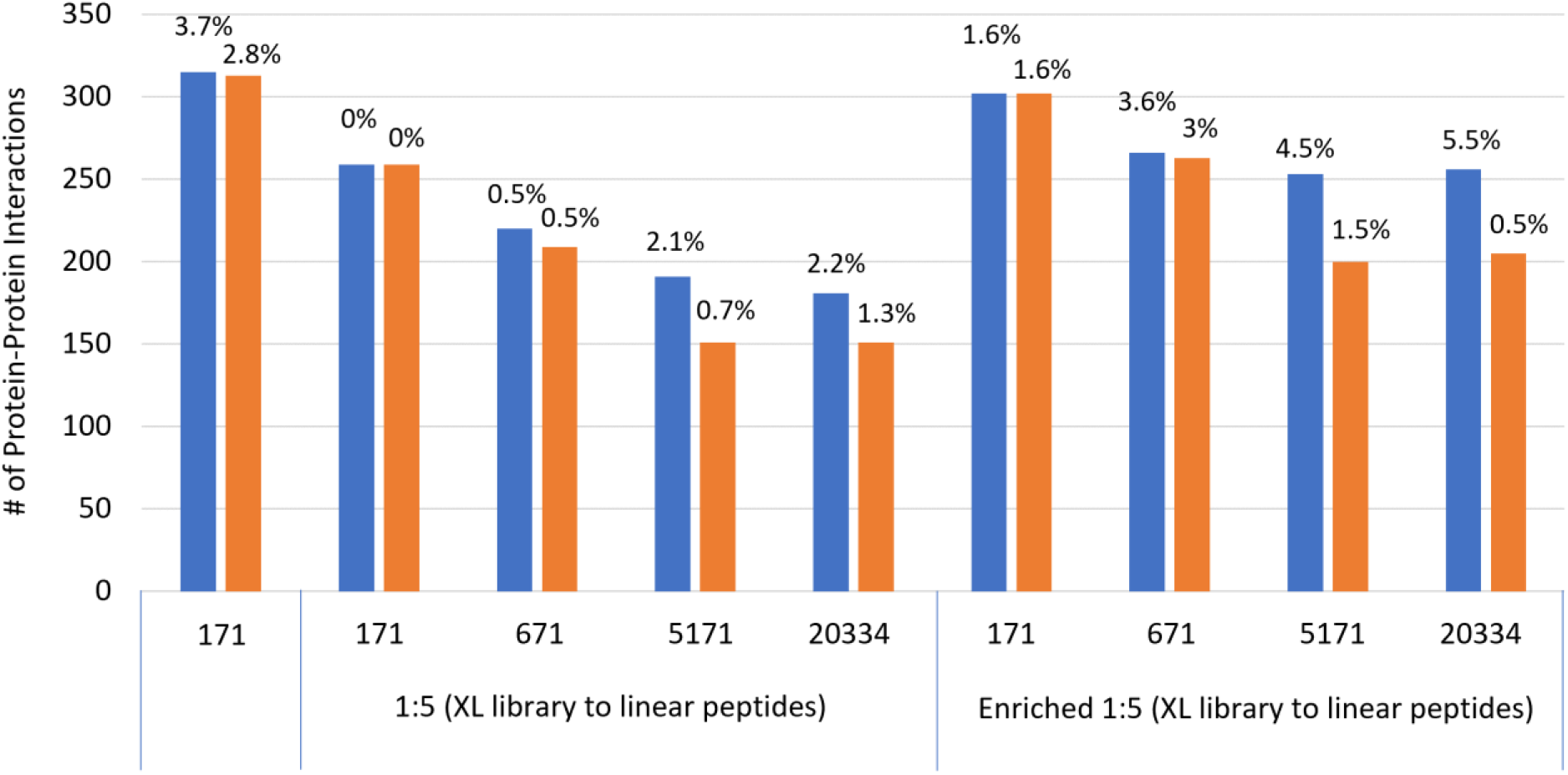
Number of detected protein-protein interaction in synthetic peptide benchmark dataset 2, from Matzinger *et al*.^30^ Blue bars represent a search conducted at an estimated 5% FDR and orange bars a search conducted at an estimated 1% FDR. All three sets of data from the benchmark were searched, with the indicated numbers of proteins in the search database to explore the effect of entrapment. Real FDR values are indicated as callouts.

However, a test on a dataset of high quality and limited complexity is not a perfect mimic of a full *in situ* crosslinking experiment. Therefore, we next analyzed the *E. coli* BS3 and DSSO crosslinking experiments from the Rappsilber group, involving 314 samples total, each analyzed with a 90 min LC-MS/MS runs on a QExactive^38^. Because no enrichment was used for this analysis beyond restricting the charge states selected during the DDA experiments, the dataset presents the full range of reaction products, both in terms of the type of reaction product and their yields. Using a single high-abundance fraction from the dataset, we conducted a grid search to determine the optimal database search settings. Interestingly, we did not detect a need to expand N beyond 10 in the library reduction phase, where we note that the setting reflects the number of unique scoring groups and not individual peptides (i.e., there can be multiple equal-scoring peptides per group). True peptides were almost always found at the highest ranks of the list, indicating that our scoring method is effective for pre-searching linearized peptides in high-complexity states. Similarly, we found that an E_α_ of 10% and an E_β_ of 99% provides optimal sensitivity, the same values used in the smaller datasets. Using these settings, all runs were searched as a single set for each crosslinker on a desktop, against the slightly restricted search space used in the original study (approximately half of the proteome)^38^. The search generated identifications for all product types, where not surprisingly the inter-protein links were detected at the lowest abundance (Table 1).

**Table 1.** Identification of reaction products from the combined search of LC-MS/MS runs from an extensively fractionated *in situ* crosslinking experiment applied to *E. coli* cells, for both BS3 and DSSO, at a 1% FDR.

|  | Inter-links | Homotypic links# | Intra-links | loop links | Mono-links | free peptides |
| --- | --- | --- | --- | --- | --- | --- |
| **BS3** | 12,415 | 3048 | 61,032 | 13,436 | 53,723 | 416,465 |
| **DSSO** | 20,104 | 5149 | 78,153 | 10,270 | 61,412 | 331,577 |

#indicates a crosslink to two identical or overlapping sequences

The CRIMP run completed in 43 hrs for BS3 (168 raw datafiles) and 39 hours for DSSO (146 raw datafiles) using modest hardware (10-core Inter i9-10850K 3.60GHz CPU with 32 GB of RAM), which is a reasonable time commitment given that each dataset required over 30 days to collect. We have found it useful to conduct an ultrafast search with E_α_ and E_β_ set to 0.01% and 25% respectively, to gauge the success of the crosslinking reaction, as such values will find high quality crosslinks. When preliminary results look promising, the samples can then be processed with greater sensitivity using the default settings.

Finally, the searches reveal a higher number of protein-protein interactions than first reported^38^. At 5% FDR, we calculate a total of 1820 PPIs for a nominal 5% FDR (compared with 756 in the original study) and 1254 PPIs at a nominal 1% FDR (compared with 590 in the original study). The calculated FDR provides a reasonable estimate of the real error rate, given the approximate nature of the true PPI space (**Figure 7**). We note that the sensitivity arises from the compositional boost, as it increases the number of hits over two-fold compared with the composition-naïve scoring method we first employed in the software. It supports the notion that the PPI search space is likely being overestimated in constructing an “all with all” interaction database. That is, adding an additional layer of information raise the scores of true interactors above a noise distribution that is likely wider than it should be. Interestingly, the overlap between the two reagents is only 20% at 5% FDR, increasing only slightly to 25% at 1% FDR, highlighting how differences in reagent design can affect sampling of the interactome.

**Figure 7.**
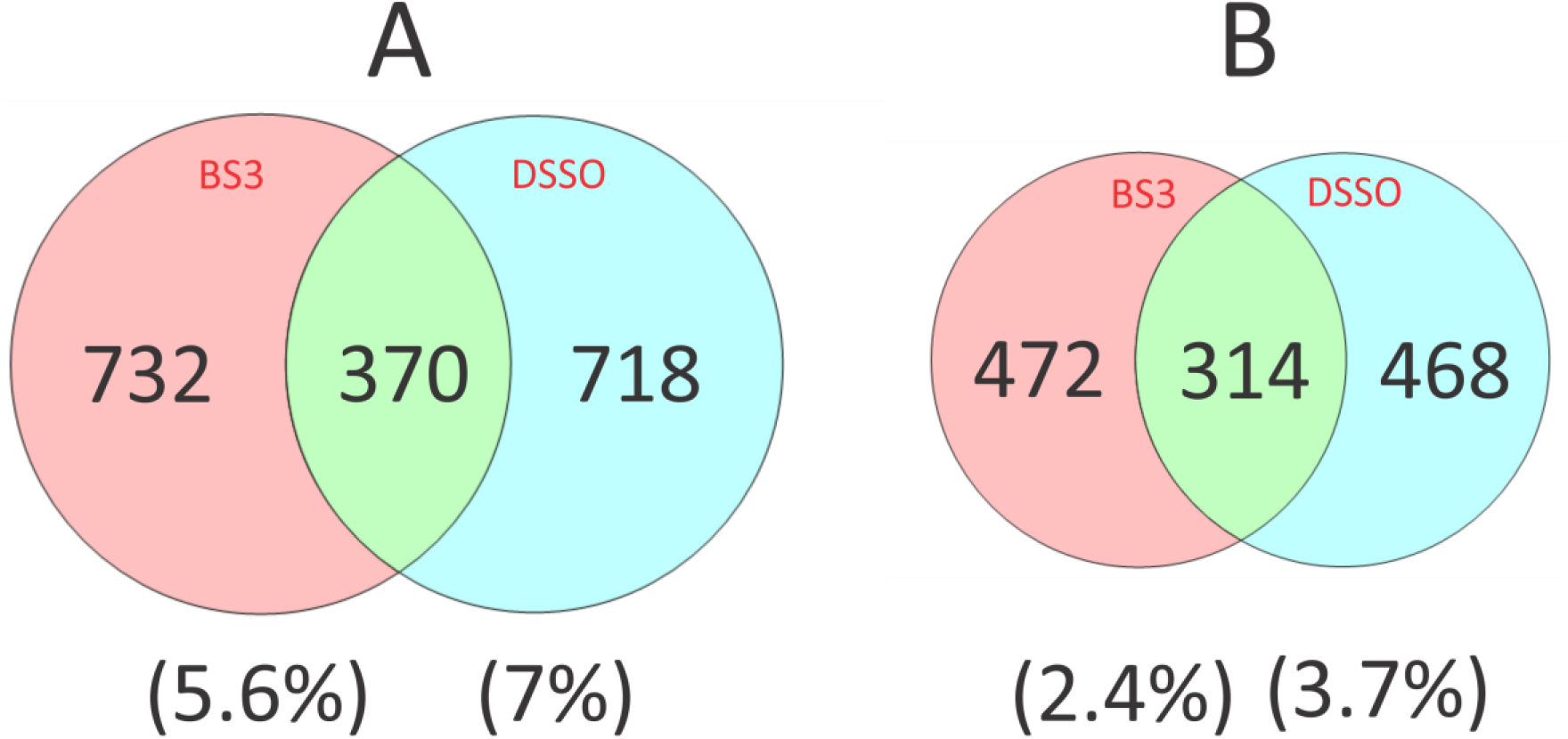
PPI search results for the *in-situ* crosslinking of the *E. coli* proteome using two crosslinking reagents. (A) searches conducted at a targeted 5% FDR and (B) searches conducted with a targeted 1% FDR. Results are based on the approximate PPI database established in Lenz *et al*.^38^, using the composition-informed PPI scoring method. Percentages at the bottom of the figure show calculated FDR values based on the composition of the library.

## Conclusions

Our results demonstrate, through the analysis of previously collected datasets, that a 2-pass database reduction method can return sensitive measurements of crosslinking composition in complex sample types. Controlling the degree of database restriction allows the user to tune the search speed to meet the needs of the experiment, without causing great concern over lost sensitivity as the dependency on the E_α_ and E_β_ scores is modest and predictable. Indeed, it is possible that brute force methods may prove to be less sensitive for highly complex systems than 2-pass methods. An unnecessary expansion of the database may generate a noisy search, much like proteomics searches do when they are parameterized with excessive numbers of variable modifications. CRIMP allows for a robust search of both cleavable and noncleavable crosslinkers alike. Noncleavable reagents should get more attention for *in situ* applications.

These reagents are easier to synthesize and are clearly complementary at this scale. Additionally, these reagents generate cross-peptide fragment ions that may be essential in validating hits, particularly when exploring highly complex states where interactions are defined by post-translational modifications. CRIMP 2.0 offers the sensitivity and search speed required for such activities.

## ASSOCIATED CONTENT

### Supporting Information

The Supporting Information is available free of charge on the ACS Publications website.

- **Additional experimental details**
  - Database segmentation for synthetic peptide benchmark dataset 1.
  - CRIMP search settings for synthetic peptide benchmark dataset 1.
  - CRIMP search settings for synthetic peptide benchmark dataset 2.
  - CRIMP search settings for proteome-scale *E. coli* dataset
- **Scoring XLs in CRIMP 2.0**
  - **Core OMSSA++ score**.
  - **Multiple Perspectives Scoring Strategy**.
  - **Calculating the Competitive Label Assignment Method (CLAM) Composite Score**
- **Error estimation in CRIMP 2.0**
- **Aggregation**
- **Composition-informed PPI scoring**
- **Figure S1**. 3D sensitivity plots for N, E_α_ and E_β_.
- **Figure S2**. A comparison of several crosslink search tools used on the synthetic peptide dataset benchmark 1, from Beveridge *et al*.^36^
- **Figure S3**. Effect of entrapment on database search times.
- **Figure S4**. Benchmarking the performance of XL search tools on complex samples, with database entrapment.
- **Figure S5**. Analysis of the DSSO crosslinking data from the synthetic peptide benchmark dataset 2, for detection of PPIs.

## AUTHOR INFORMATION

### Author contributions

DAC and VS were responsible for software design and coding. DAC, VS and DCS contributed to algorithm design. The software was extensively evaluated by all authors. DAC, VS and DCS wrote the final manuscript with input and editing from the other coauthors.

### Notes

The authors declare no competing financial interest.

## ACKNOWLEDGMENT

The authors would like to thank the many members of the community for extensive feedback provided on the use of CRIMP 2.0 for crosslinking applications. This work was funded by the Natural Sciences and Engineering Research Council of Canada Discovery Grant (RGPIN 2017-04879 and CANARIE grant RS3-084.

