## Supporting Information for "High sensitivity proteome-scale search for crosslinked peptides using CRIMP 2.0"

**Contents:**


1. **Additional experimental details**
    a. Database segmentation for synthetic peptide benchmark dataset 1.
    b. CRIMP search settings for synthetic peptide benchmark dataset 1.
    c. CRIMP search settings for synthetic peptide benchmark dataset 2.
    d. CRIMP search settings for proteome-scale *E. coli* dataset
2. **Scoring XLs in CRIMP 2.0**
    a. **Core OMSSA++ score.**
    b. **Multiple Perspectives Scoring Strategy.**
    c. **Calculating the Competitive Label Assignment Method (CLAM) Composite Score**
3. **Error estimation in CRIMP 2.0**
4. **Aggregation**
5. **Composition-informed PPI scoring**
6. **Figure S1.** 3D sensitivity plots for N, $E_\alpha$ and $E_\beta$.
7. **Figure S2.** A comparison of several crosslink search tools used on the synthetic peptide dataset benchmark 1, from Beveridge *et al.*[1]
8. **Figure S3.** Effect of entrapment on database search times.
9. **Figure S4.** Benchmarking the performance of XL search tools on complex samples, with database entrapment.
10. **Figure S5.** Analysis of the DSSO crosslinking data from the synthetic peptide benchmark dataset 2, for detection of PPIs.

1. **Additional experimental detail**
   a. **Database segmentation for synthetic peptide benchmark dataset 1.**

>Cas9a
GAASMDKKYSIGLAIGTNSVGWAVITDEYKVPSKKFKVLGNTDRHSIKKNLIG
ALLFDSGETAEATRLKRTARRRYTRRKNRICYLQEIFSNEMAKVDDSFFHRLE
ESFLVEEDKKHERHPIFGNIVDEVAYHEKYPTIYHLRKKLVDSTDKADLRLIYL
ALAHMIKFRGHFLIEGDLNPDNSDVDKLFIQLVQTYNQLFEENPINASGVDA
KAILSARLSKSRRLENLIAQLPGEKKNGLFGNLIALSLGLTPNFKSNFDLAE
DAKLQLSKDTYDDDLDNLLAQIGDQYADLFLAAKNLSDAILLSDILRVNTEITKA
PLSASMIKRYDEHHQDLTLLKALVRQQLPEKYKEIFFDQSKNGYAGYIDGG
ASQEEFYKFIKPILEKMDGTEELLVKLNREDLLRKQRTFD

>Cas9b
NGSIPHQIHLGELHAILRRQEDFYPFLKDNREKIEKILTFRIPYYVGPLARG
NSRFAWMTRKSEETITPWNFEEVVDKGASAQSFIERMTNFDKNLPNEKVLPK
HSLLYEYFTVYNELTKVKYVTEGMRKPAFLSGEQKKAIVDLLFKTNRKVTVK
QLKEDYFKKIECFDSVEISGVEDRFNASLGTYHDLLKIIKDKDFLDNEENED
ILEDIVLTLTLFEDREMIEERLKTYAHLFDDKVMKQLKRRRYTGWGRLSRKL
INGIRDKQSGKTILDFLKSDGFANRNFMQLIHDDSLTFKEDIQKAQVSGQGD
SLHEHIANLAGSPAIKKGILQTVKVVDELVKVMGRHKPENIVIEMARENQTT
QKGQKNSRERMKRIEEGIKELGSQILKEHPVEN

>Cas9c
TQLQNEKLYLYYLQNGRDMYVDQELDINRLSDYDVDAIVPQSFLKDDSIDNK
VLTRSDKNRGKSDNVPSEEVVKKMKNYWRQLLNAKLITQRKFDNLTKAERGG
LSELDKAGFIKRQLVETRQITKHVAQILDSRMNTKYDENDKLIREVKVITLK
SKLVSDFRKDFQFYKVREINNYHHAHDAYLNAVVGTALIKKYPKLESEFVYG
DYKVYDVRKMIAKSEQEIGKATAKYFFYSNIMNFFKTEI

>Cas9d
TLANGEIRKRPLIETNGETGEIVWDKGRDFATVRKVLSMPQVNIVKKTEVQT
GGFSKESILPKRNSDKLIARKKDWDPKKYGGFDSPTVAYSVLVVAKVEKGKS
KKLKSVKELLGITIMERSSFEKNPIDFLEAKGYKEVKKDLIIKLPKYSLFEL
ENGRKRMLASAGELQKGNELALPSKYVNFLYLASHYEKLKGSPEDNEQKQLF
VEQHKHYLDEIIEQISEFSKRVILADANLDKVLSAYNKHRDKPIREQAENII
HLFTLTNLGAPAAFKYFDTTIDRKQYRSTKEVLDATLIHQSITGLYETRIDL
SQLGGD

#### b. CRIMP search settings

| Parameters changed | Synthetic Benchmark 1 | | Synthetic Benchmark 2 | Proteome Search | |
|---|---|---|---|---|---|
| **Crosslinker** | DSS | DSSO | DSSO | BS3 | DSSO |
| **Digest** | Trypsin (K/R only) | Trypsin (K/R only) | Trypsin (K/R only) | Trypsin (K/R only) | Trypsin (K/R only) |
| **Eα** | 99% | 99% | 99% | 99% | 99% |
| **Eβ** | 25% | 25% | 10% | 10% | 10% |
| **Top N** | 10 | 10 | 10 | 10 | 10 |
| **Min. Peptide Size** | 5 | 5 | 6 | 6 | 6 |
| **Max. Peptide Size** | 60 | 60 | 30 | 30 | 60 |
| **Min. m/z Range** | 400 | 400 | 375 | 400 | 400 |
| **Min. m/z Range** | 1600 | 1600 | 1500 | 1450 | 1450 |
| **Database Propagation** | Dataset | None | None | None | Run/State Group |
| **MS1 Tolerance** | 5ppm | 5ppm | 5ppm | 5ppm | 5ppm |
| **MS2 Tolerance** | 20ppm | 20ppm | 10ppm | 10ppm | 10ppm |
| **Fragmentation Events** | 2 | 1 | 1 | 2 | 1 |
| **Exclude Y1s (Processor)** | FALSE | FALSE | TRUE | TRUE | TRUE |

**Table S1. A list of search parameters for each dataset.** The parameters highlight modified (non-default) parameters values plus the default TopN, Eα and Eβ settings, and follow the specifications of each benchmark publication as close as possible.

### 2. Scoring XLs in CRIMP 2.0

**a. Core OMSSA++ score.** As described in the main text, we revised our peak identification process to better support a revised scoring method. The essence of the scoring method remains the OMSSA+ score defined in Sarpe *et al.*[2], which assigns a probabilistic E-value based loosely on the original OMSSA concept[3]. Here, in the $MS^1$ space, our new peak identification process improves our use of the precursor ion isotopic envelope and changes the way in which precursor mass is handled. In $MS^2$ space, fragments are assigned using a "greedy feature overlap solver (GFOS)", that assigns weights, or priorities, to fragment types in the $MS^2$ spectrum based on a hierarchical approach to spectral assignment. GFOS also partially deconvolves overlapped isotopic distributions, reduces the frequency of monoisotopic misassignments and improves charge identification. We refer to the collection of enhancements as generating a core **OMSSA++ score** for a given $MS^2$ spectrum. We note also that annotation support and scoring is provided for cleavable reagents and their fragment types, based on doublet detection for the component peptides as well as their fragments.

**b. Multiple Perspectives Scoring Strategy.** The OMSSA++ score, or any single scoring method, does not readily support distinguishing between hits that draw from variable database sizes, as occurs when considering the different reaction products arising from a crosslinking reaction. The various fragment categories that can arise from these reaction products also contain different information about spectrum matches. For example, crosslink-specific fragments will indicate a crosslink has occurred but these are of less value in determining the sequences that comprise the crosslink. Finding a best solution (here, the best-scoring reaction product) can be approached using strategies derived from computer vision applications (such as autonomous vehicle guidance). The spectral identification process is like viewing an image from multiple perspectives, where each viewpoint captures overlapping sets of information, and synthesizes the information to classify an object.

We developed a **Multiple Perspectives Scoring (MSP**) process to generate better spectral "depth perception", involving the creation of a multi-component scoring vector with the following elements, each scored separately using OMSSA++:

- All annotated single-fragmentation events.
- All annotated fragments from a linear free (*i.e.*, noncrosslinked) peptide.
- All annotated "single-fragment" crosslinks, which refers to a single sequence fragment in a crosslinked peptide (*i.e.*, with retention of the linkage).
- All annotated fragments that support the most likely crosslinked residue pair, including internal (or two-fragment) ions.
- All annotated fragments, including internal ions.

These components were selected to represent fragment classes differently in the final score, preventing the noise from potentially large classes of fragments (such as internal ions) from overwhelming other low-complexity annotations (such as single-fragment crosslinks). The fragment allocation process used in OMSSA++ scoring is synergistic with this strategy. The scoring vector is augmented with an equivalent set of scores for the weakest of the two component peptides, creating 10 probabilistic components. These are further augmented with an additional 4 components that contribute non-probabilistic (and class-independent) spectral information, to create a 14-component scoring perspective.

**c. Calculating the Competitive Label Assignment Method (CLAM) Composite Score.** We then score the $MS^2$ spectra using a three-stage process:

- First, spectral matches are scored independently for all reaction products, assuming no spectral conflicts, using the 14-component vector.
- Second, spectral conflicts are identified using only the probabilistic components of the soring vector. All matches competing for a spectrum are evaluated in a "one vs. rest" strategy to generate a γ discriminator for a given spectral assignment, *j*:

$$\gamma_j = n \times \frac{x_j}{1 + max\,\{x_1 \dots x_n\}}$$

where *j* is an element in a vector of all *n* spectral conflicts and

$$x_j = \sum_i^k \left(-\ln(E_{total,i}) + \ln(E_{shared,i})\right)_j$$

$E_{total,i}$ is the OMSSA++ score for the $i^{th}$ component of the score vector for the full set of fragments for a given spectral assignment, and $E_{shared,i}$ is the OMSSA++ score for the $i^{th}$ component of the vector for the shared set of fragments arising from all other possible spectral assignments; k is number of components in the scoring vector.

Then, the total fragment set for a given spectral assignment is penalized by the cumulative shared fragment set from other possible assignments. A $\delta_i$ score is calculated for the given assignment as follows:

$$\delta_i = -\ln(E_{total,i}) + \ln\left(\frac{\gamma_j}{1-\gamma_j} E_{shared,i}\right)$$

where $\delta_i$ is a penalized score for the given vector component $i$, upon which point the non-probabilistic components of the vector are reintroduced.

- Third, each term in the scoring vector is then scaled (0,100) and marginal q values are calculated for each component (using the decoy information and FDR estimation described below) and applied to the δ score:

$$D_i = (1 - q_i)\delta_i$$

This transforms the scores vector by the specific error function of each component. Finally, a CLAM score is calculated, reducing the vector to a scalar quantity, through the log-link of the inner product:

$$CLAM\ score = \ln\left(\sum_i^k \sum_j^k D_i D_j\right)$$

In summary, the MSP scoring approach generates a CLAM score that identifies the best hit across all categories of reaction product. The final CLAM score is rescaled (0,100) for ease of "in run" comparisons. They are not directly comparable between experiments.

### 3. Error estimation in CRIMP 2.0

The different categories of crosslinker reaction products all have different noise distributions in database searches, leading most tools to calculate False Discovery Rates (FDRs) in a category-specific manner. The strategy adopted in CRIMP 2.0 removes the need for category-specific error estimation. First, database matches are grouped according to their unique database configurations:

- Free linear peptides
- Mono-link peptides
- Loop-linked peptides
- Intra-protein crosslinking peptides
- homotypic inter-protein crosslinks

- heterotypic inter-protein crosslinks.

Within each of these groups, the match scores are standardized by their error distributions (approximated with a Poisson mixture model, to account for uneven sampling of decoys). This establishes a situation where the FDR category-specific error distributions are essentially a sampling of a universal error distribution, which allows us to directly combine the results across the categories and achieve a more robust single error estimate. Only then are the scores scaled (0,100). Next, we calculate both a global and a local q value and combine them in a geometric mean as follows, to achieve a category-independent FDR estimate:

- Global q-estimate

$$q_{global} = \frac{(N_D + N_{TD}) - N_{DD}}{(N_T + N_{TT})} \in \{ Decoys \mid Score \geq T_i \}$$

- Local/Tailed q-estimate

$$q_{local} = \frac{(N_D + N_{TD}) - N_{DD}}{(N_T + N_{TT})} \in \begin{cases} (Decoys \mid Score \geq T_j) \\ (Targets \mid Score \geq T_j) \end{cases}$$

- Final estimate

$$FDR = \sqrt{q_{global} * q_{local}}$$

where $N_D$ is the number of decoy hits for a peptide, $N_T$ the number of target hits for a peptide, $N_{TD}$ the number of target-decoy hits for a crosslink, $N_{DD}$ the number of decoy-decoy hits, and $N_{TT}$ the number of target-target hits.

**4. Aggregation**

To propagate spectral matches into higher levels of information, we need to implement a score cut-off to determine the set of significant matches to use as inputs for aggregation. Here the score cut-off is calculated by minimizing the equal-error rate (EER, also known as the crossover-error rate (CER)). Matches with scores greater than the EER value are accepted and passed forward to aggregation. The EER score is the smallest score that minimizes the distance between the FDR and the FOR (false omission rate) and is calculated as follows:

*A. EER point estimate*

$$EER = FDR - FOR$$

B. *EER score cut-off (smallest root of EER function)*

$$\alpha_{score} = \min_{EER \to 0}(score)$$

This approach avoids artificially truncating positive results prior to the scoring aggregation process, which translates into greater sensitivity and a more accurate error estimation for the higher levels of information.

Results are then aggregated within a run and, if replicate samples are processed, across multiple runs. CSMs are propagated to higher levels of information (*e.g.*, UPPs) using the following equation:

$$CLAM_{i+1} = \frac{max(CLAM_i) + \sum_i^n CLAM_i}{n+2}$$

Where subscript $i$ represents the information level (e.g., CSMs is an $i = 1$). At the conclusion of each aggregation step, scores are rescaled and final FDR calculations are applied as above for each level of information.

### 5. Composition-informed PPI scoring

It has been noted elsewhere and confirmed by our own experience that the true set of PPIs should have strong evidence for their unique proteins in addition to the PPI specific evidence. Our composition-informed PPI score includes information from the unique proteins in a three-component vector consisting of:

- AB-Score, composed of inter-protein dimers, and calculated as above through aggregation.
- A-Protein Score, composed of intra-protein crosslinks + monomers and calculated as above through aggregation.
- B-Protein Score, composed of intra-protein + monomers and calculated as above through aggregation.

The 3-component scores vector is collapsed to a single scalar using an adaptation of our scoring workflow. Here, the marginal q-value transformation and inner-product scoring method of the CLAM process are reapplied to the set of component scores. The recalculated score is calculated as follows:

- *Marginal q transformation applied to PPI score vector*

$$A_i = (1 - q_i) \times \alpha_i$$

- *Final PPI score calculation, adapted CLAM product*

$$PPI\ Score = (A_{PPI} \times A_{PPI}) + (A_{PPI} \times A_{PA}) + (A_{PPI} \times A_{PB}) + (A_{PA} \times A_{PB})$$

After the score recalculation, a final FDR estimate is obtained, and the PPI q-values are replaced by the new estimate directly. Like the other scores, the final recalculated PPI scores are min/max scaled to [0,100] where 100 indicates the most confident match score.

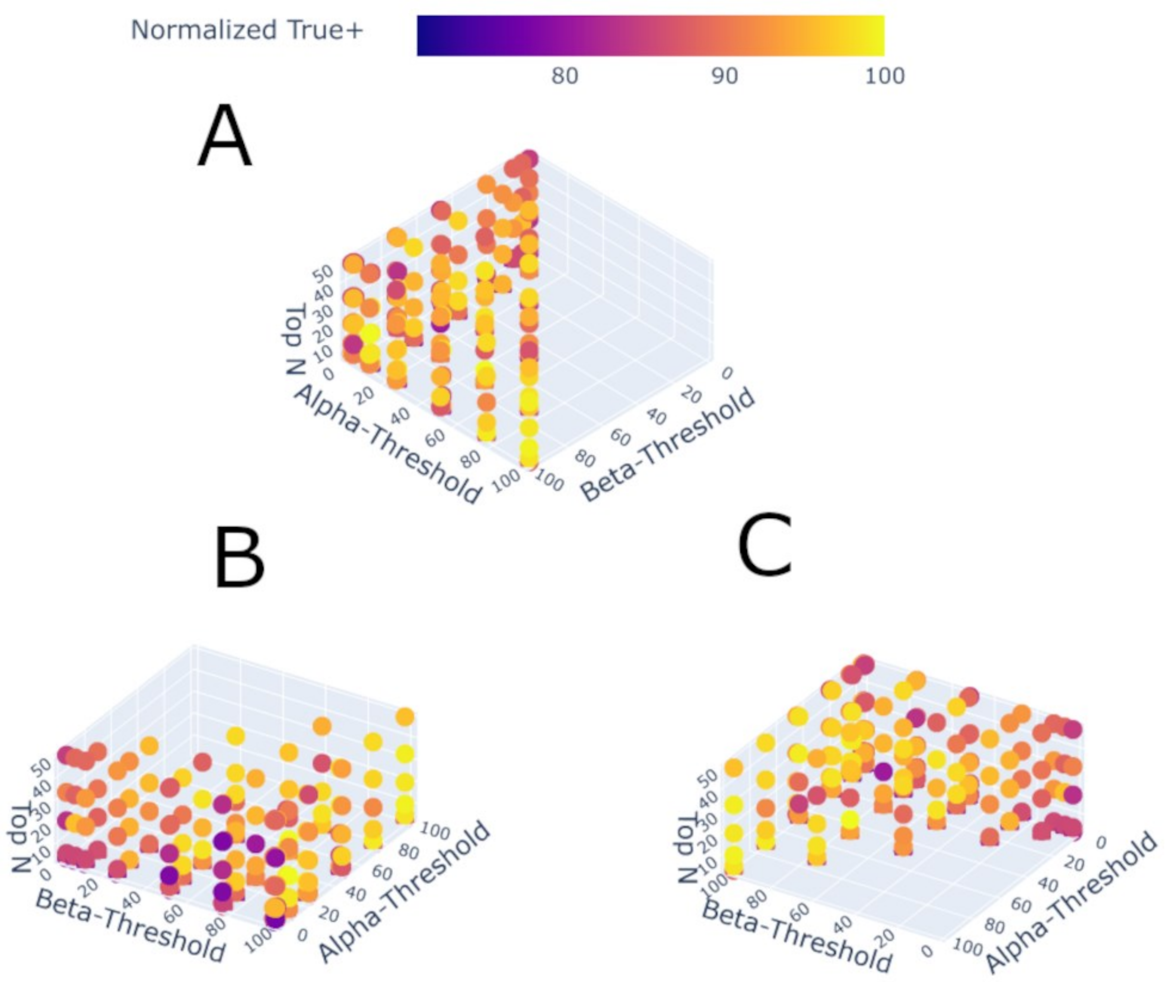


**Figure S1. 3D Sensitivity plots for N, Eα and Eβ, using DSS data from Beveridge *et al.*[1] dataset.** A grid search of the library reduction parameter space was conducted with increasing entrapment complexity up to addition of the *E. coli* proteome. Sensitivity was normalized per entrapment database per run to facilitate comparison. Search settings used for primary experiments (Eb=99, Ea=25, N=10) was chosen for its balance between search time, and sensitivity (mean=95.37%, min=84.15%, max=98.63%, Q1=95.84%, Q2=97.06%, Q3=98.1%), n=24).

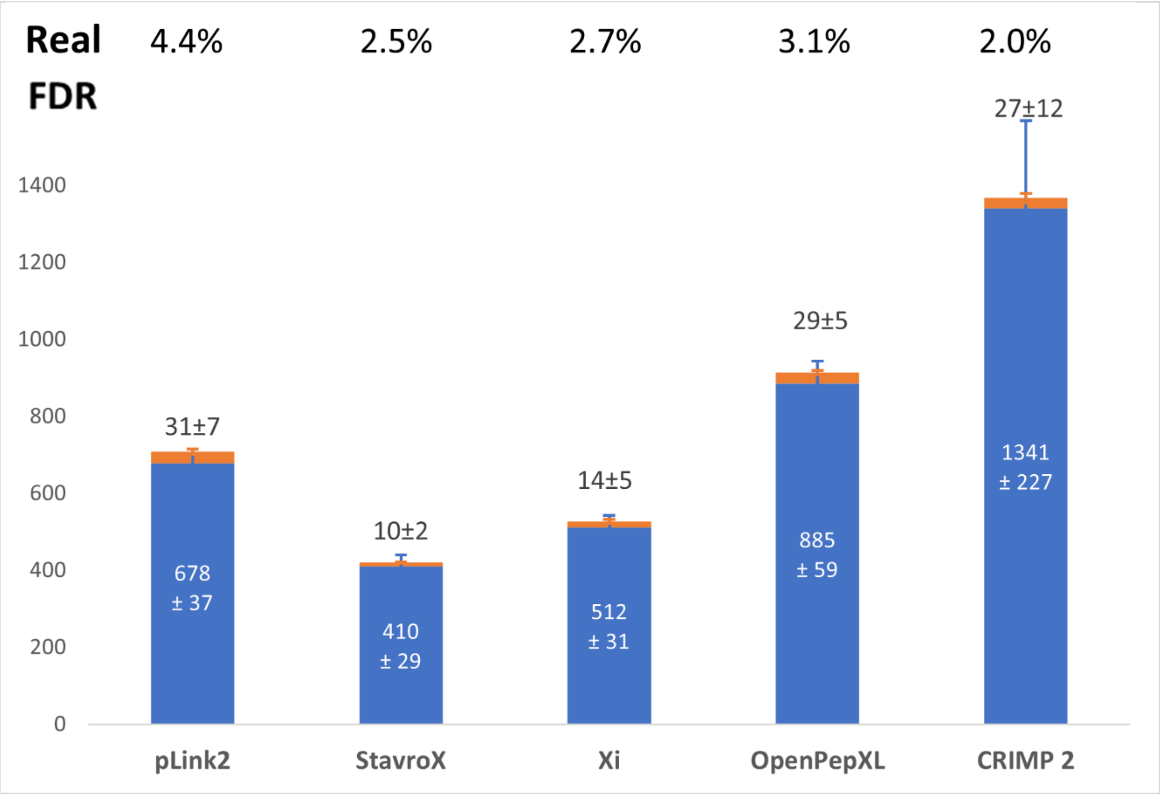


**Figure S2. A comparison of several crosslink search tools used on the synthetic peptide dataset benchmark 1, from Beveridge *et al.*[1]** The number of CSMs that correspond to correct (blue) and incorrect (orange) crosslinks identified by the indicated search tools. Results were filtered to an estimated 5% FDR at the CSM level, and the real FDR, based on knowledge of the composition of the dataset (the real FDR) is indicated. Data were searched with 10 additional proteins as entrapment. Error bars correspond to the standard deviation from technical replicates (n=3).

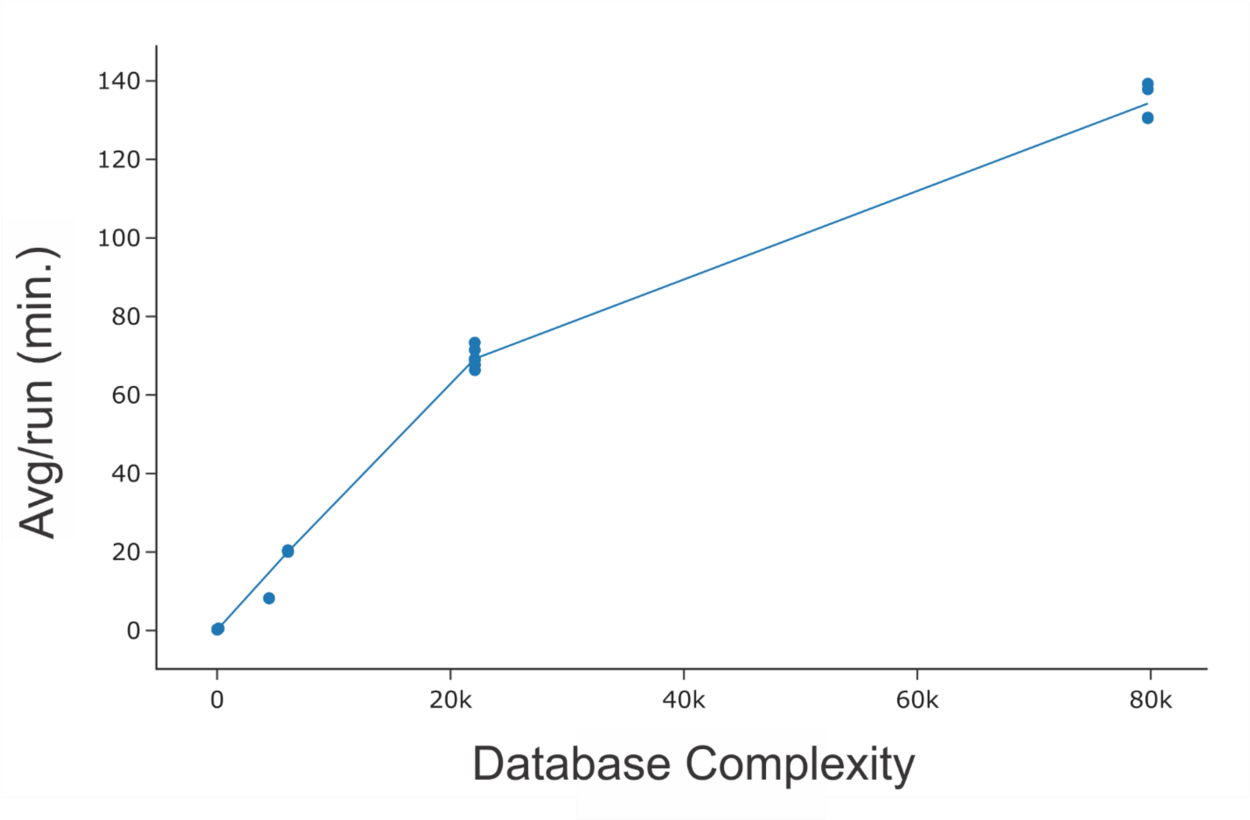


**Figure S3. Effect of entrapment on database search times**. Search used an AMD Ryzen 7 - 5800X computer (16-logical processors, 3.8 GHz, 32 GB RAM), operating on the synthetic peptide benchmark dataset 1, from Beveridge *et al.*[1]

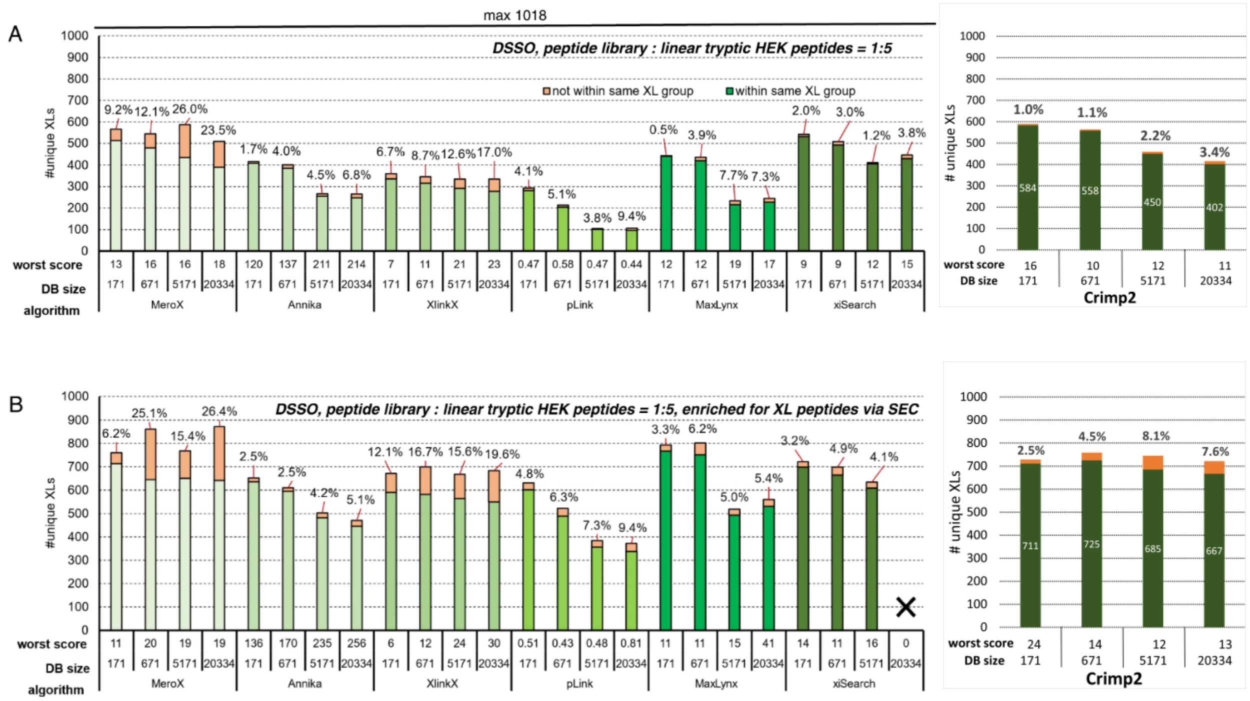


**Figure S4**. **Benchmarking the performance of XL search tools on complex samples, with database entrapment.** (A) Direct measurement of XLs from the DSSO-linked library mixed with linear tryptic HEK 293 digest peptides (1:5 w/w) (B) As in A, but after enrichment with size exclusion chromatography. Bars indicate the number of unique crosslinks identified using the indicated algorithm at a 1% estimated FDR for databases of 171 *E. coli* ribosomal proteins, or additional human proteins as noted. Green-spectrum bars show true positives, and orange bars show false positives. Callouts show actual FDR, and the lowest score at the FDR cutoff is also shown for each algorithm. Figure reproduced from Matzinger et al.[4] (with permission) with the addition of Crimp2 search data from this study.

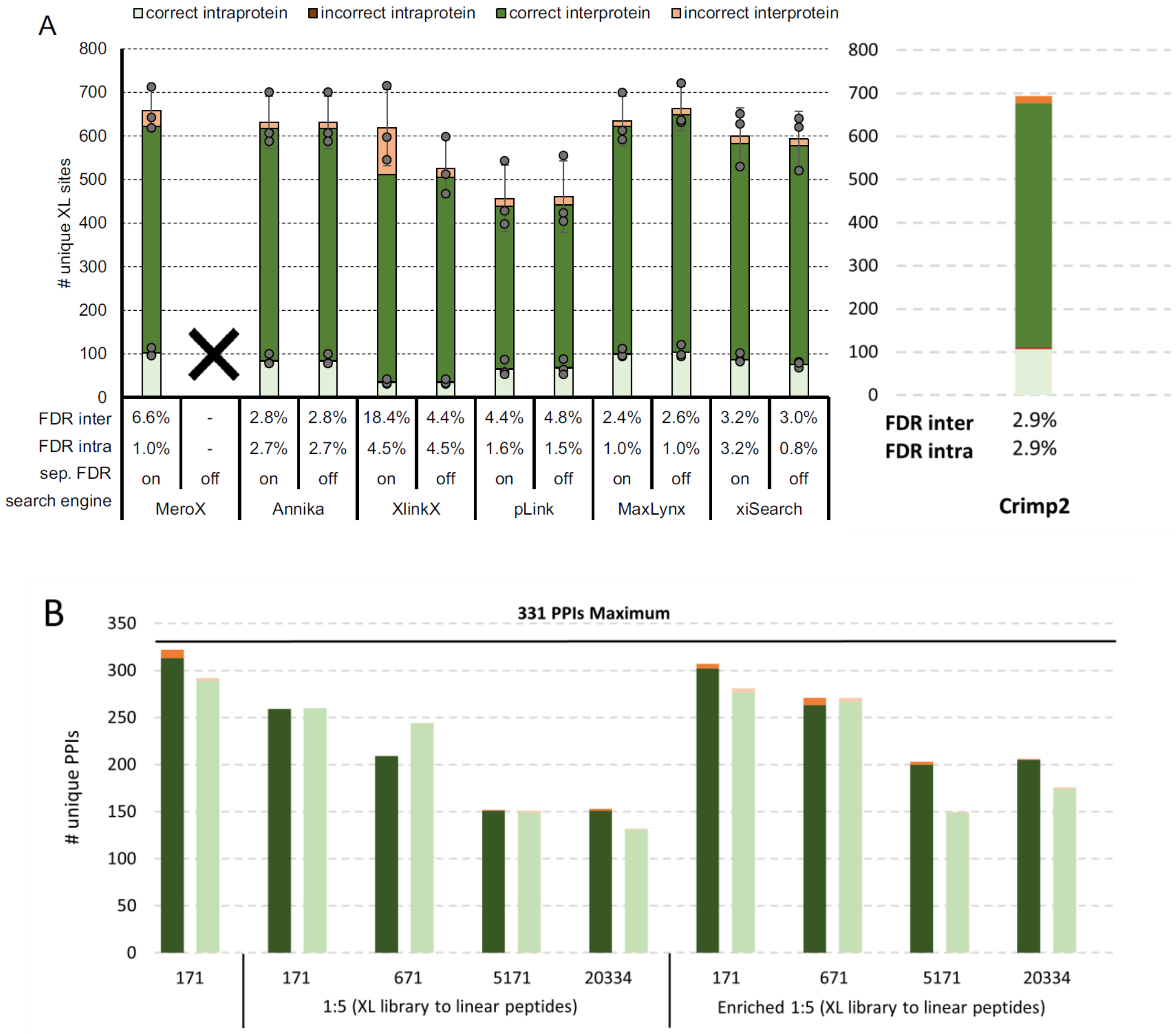


**Figure S5. Analysis of the DSSO crosslinking data from the synthetic peptide benchmark dataset 2, for detection of PPIs** (A) The enumeration of inter-protein and intra-protein crosslinks at a nominal 1% FDR from the 171-protein database search, as a function of algorithm type and allowing for the calculation of separate or combined FDR calculations where possible. Figure reproduced from Matzinger *et al.*[4], with the addition of a CRIMP 2.0 analysis. (B) PPI analysis of the same data, with the addition of the peptide-contaminated data and the enriched data, as noted. Data searched at a nominal FDR of 1%, with the indicated database size. Dark green and orange bars show the true and false positives (respectively) for the composition-informed PPI scoring method. Light green and orange bars show the true and false positives (respectively) for a composition-naïve PPI scoring method, where the score is based solely on interprotein crosslinks.
